# Polygenic Link Between Blood Lipids And Amyotrophic Lateral Sclerosis

**DOI:** 10.1101/138156

**Authors:** Xu Chen, Solmaz Yazdani, Fredrik Piehl, Patrik K.E. Magnusson, Fang Fang

## Abstract

Dyslipidemia is common among patients with amyotrophic lateral sclerosis (ALS). We aimed to test the association and causality between blood lipids and ALS, using polygenic analyses on the summary results of genome-wide association studies. Polygenic risk scores (PRS) based on low-density lipoprotein cholesterol (LDL-C) and total cholesterol (TC) risk alleles were significantly associated with a higher risk of ALS. Using single nucleotide polymorphisms (SNPs) specifically associated with LDL-C and TC as the instrumental variables, statistically significant causal effects of LDL-C and TC on ALS risk were identified in Mendelian randomization analysis. No significant association was noted between PRS based on triglycerides or high-density lipoprotein cholesterol risk alleles and ALS, and the PRS based on ALS risk alleles were not associated with any studied lipids. This study supports that high levels of LDL-C and TC are risk factors for ALS, and it also suggests a causal relationship of LDL-C and TC to ALS.

## Introduction

Dyslipidemia is common among patients with amyotrophic lateral sclerosis (ALS). Higher-than-expected levels of low-density lipoprotein cholesterol (LDL-C), total cholesterol (TC), triglycerides (TG) and low/high-density lipoprotein cholesterol ratio (LDL-C/HDL-C) have been reported among patients with ALS by many although not all studies.^1–6^ In a recent observational study, we found that patients with ALS had increased levels of LDL-C and apolipoprotein B up to 20 years before diagnosis.^7^ Whether or not there is a genetic contribution to the noted dyslipidemia in ALS is unknown, and it is also unclear whether dyslipidemia plays a causal role in ALS.

By using single nucleotide polymorphisms (SNPs) from large-scale genome-wide association studies (GWAS), polygenic methods have been developed to investigate the genetic etiology and causality between human complex traits.^8^ SNPs derived polygenic risk scores (PRS) of lipids have for instance been shown to be predictive of circulating lipid trajectories.^9,10^ Mendelian randomization (MR) study using SNPs that are specifically associated with LDL-C and TG (but not with other risk factors or confounders) has further clearly suggested causal relationships of LDL-C and TG to coronary artery diseases (CAD).^11,12^ In this study, we aimed to test the association and causality between blood lipids and ALS, by using polygenic methods on the latest GWAS results among Europeans.

## Materials and Methods

### Data

All data used in this study were summary GWAS results in European ancestry populations. Because the association and causality between blood lipids and CAD have been established by polygenic methods in previous study, we used CAD as a “point of reference” outcome to verify the data quality and parameter setting of the present analysis on lipids and ALS.

GWAS results on blood lipids were accessed from the Global Lipids Genetics Consortium, including ∼100 000 individuals (N_LDL-C_=95 454, N_TC_=100 184, N_TG_=96 598, N_HDL-C_=99 900) and 2.69 million SNPs.^13^ GWAS results on ALS were accessed from the Project MinE GWAS Consortium, including 36 052 individuals (N_case_= 12 577) and 8.71 million SNPs.^14^ GWAS results on CAD were accessed from the CARDIoGRAMplusC4D Consortium, including 86 995 individuals (N_case_=22 233) and 2.42 million SNPs.^15^

### Polygenic analyses

Bi-directional PRS analyses were performed to check the association between blood lipids and ALS, in which blood lipids were used first as base (exposure) and then as target (outcome). In the base data, correlated SNPs were clumped by using HapMap_ceu_all genotypes (release 22, 60 individuals, 3.96 million SNPs) as the linkage disequilibrium (LD) reference, with the recommended parameter settings in PRSice v1.25: including a clumping threshold of p1=p2=0.5, a LD threshold of r^2^=0.05, and a distance threshold of 300Kb; independent SNPs were then included in quantiles with gradually increasing P-value thresholds (P_T_, ranging from 0 to 0.5, in steps of 1x10^−4^ per quantile); the P_T_ of the quantile that explained the largest variance of the target was defined as the best-fitted P_T_.^16^

When a significant association was identified between certain lipid and ALS in the PRS analyses, we conducted a MR analysis to test the potential causality.^17^ PhenoScanner database was used to identify the specific loci for the lipid in question: the lead SNP in each locus achieving genome-wide significance (P-value < 5x10^−8^) and are not associated with other lipids or potential confounders of the lipid-ALS relationship (e.g., lifestyle factors and body mass index, P-value>5x10^−8^).^18^ The lead SNPs (or their proxy SNPs) in the selected loci were then used as instrumental variables (Fig. 1), and the causal effect of the exposure (lipid) on outcome (ALS) was calculated and plotted by using the “gtx v0.0.8” package in R 3.2.5.

**Fig.1.**
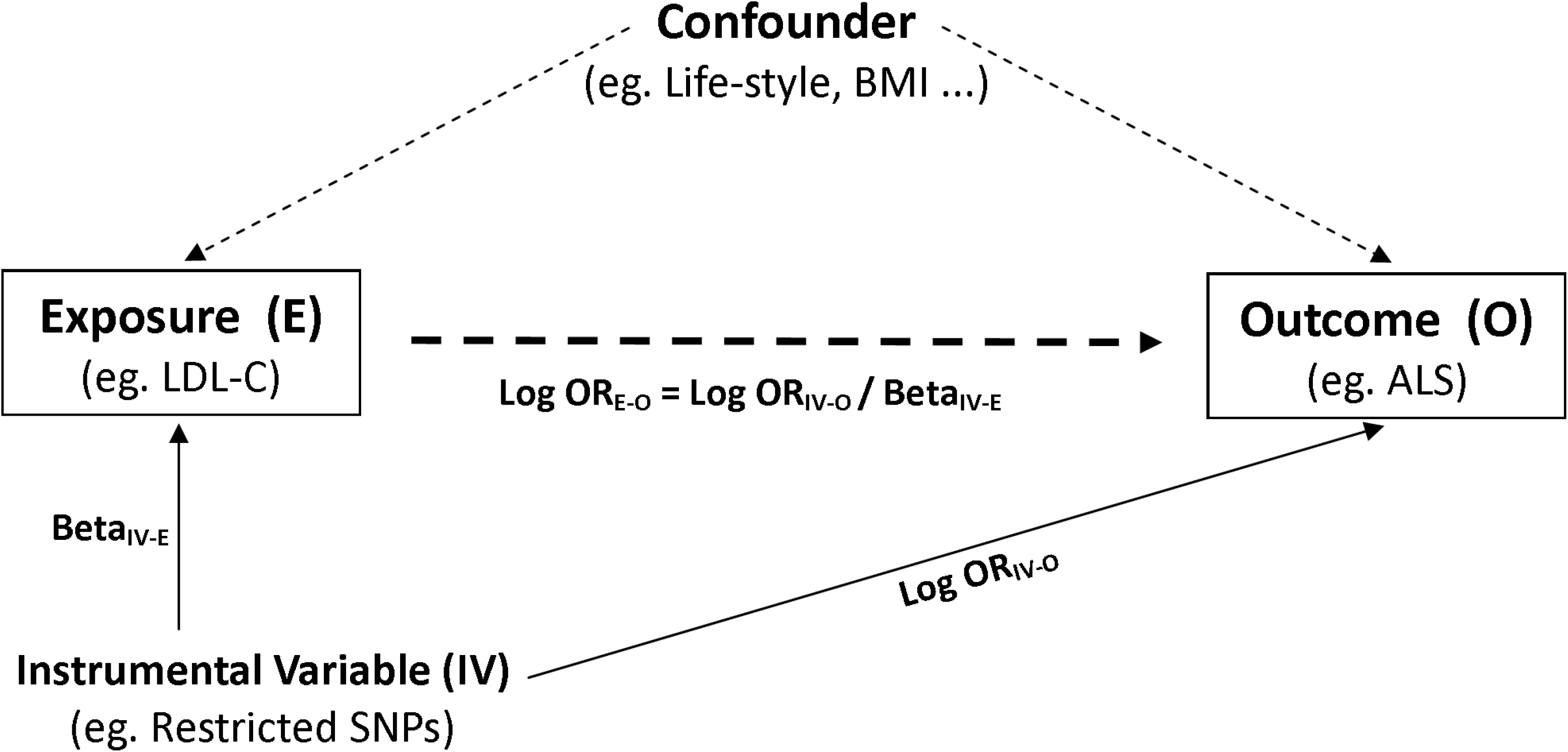
Framework of the Mendelian Randomization Analysis. Restricted single nucleotide polymorphisms (SNPs) were used as instrumental variables to test the causal effect of exposure on the outcome. OR: odds ratio; LDL-C: low-density lipoprotein cholesterol; BMI: body mass index; ALS: amyotrophic lateral sclerosis.

## Results

### Bi-directional PRS analyses

The best-fitted PRS based on LDL-C and TC risk alleles were both statistically significantly associated with ALS risk (Table 1). The odds ratios (ORs) were 1.16 (95% confidence interval [CI] 1.10-1.22, P=8.82x10-^8^) and 1.15 (95% CI 1.09-1.21, P=8.17x10-^7^) per standard deviation of the PRS based on LDL-C and TC risk alleles, respectively. The best-fitted P_T_ was the same (5x10^−5^) for LDL-C and TC, with SNPs in the corresponding quantiles explaining 0.08% and 0.07% of the variance of ALS. The associations of PRS based on LDL-C and TC risk alleles with ALS were similar and statistically significant across quantiles with P_T_⩽5x10^−5^ (Fig. 2A-B). Similar results were noted for the quantiles when including only genome-wide significance SNPs (P_T_⩽5x10^−8^), and the variance of ALS explained by these SNPs was similar to the variance explained by the best-fitted quantile (Fig. 2C-D).

**Fig.2.**
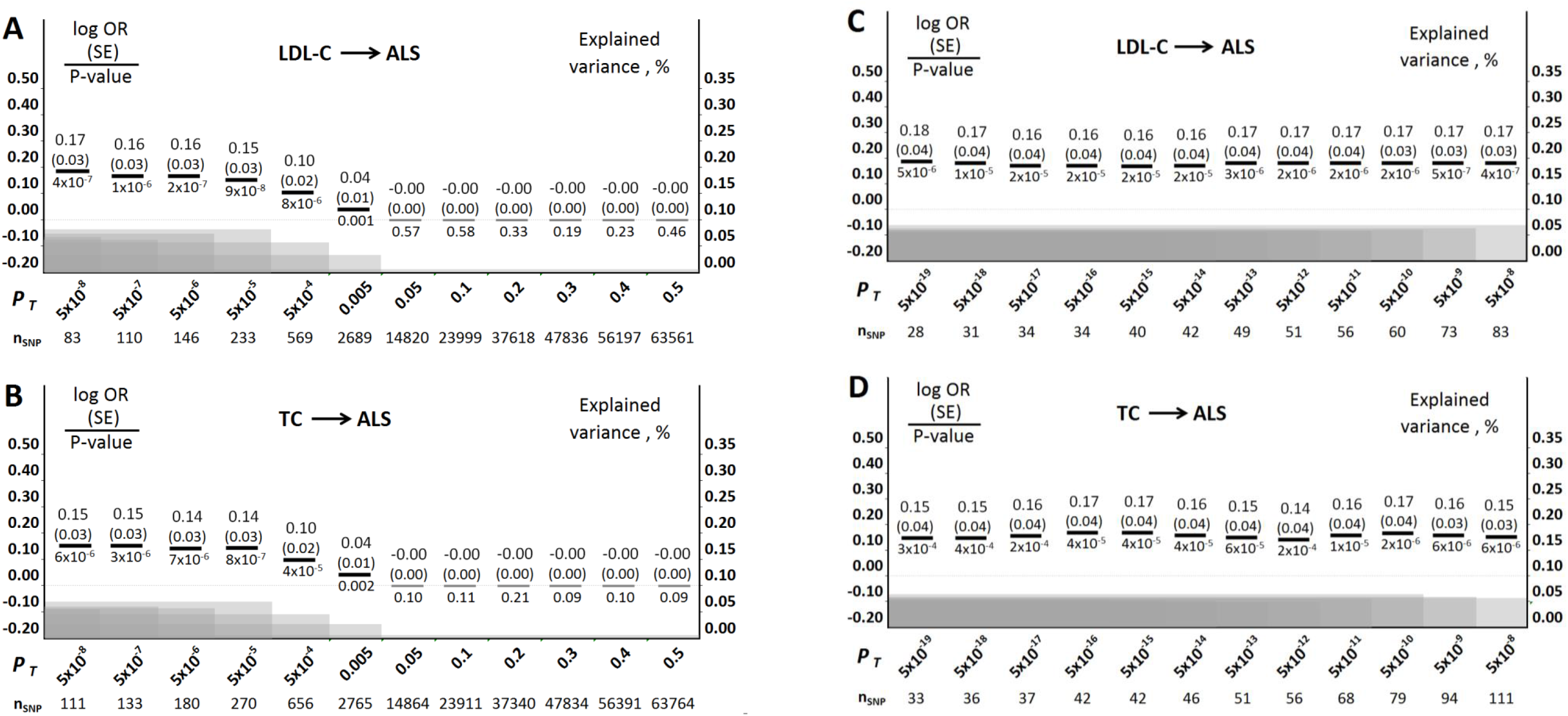
Polygenic Risk Score (PRS) Analyses of LDL-C and TC on ALS in Europeans. **(A)** PRS based on risk alleles of LDL-C (independent SNPs across the whole genome) in association with ALS. **(B)** PRS based on risk alleles of TC (independent SNPs across the whole genome) in association with ALS. **(C)** PRS based on risk alleles of LDL-C (independent SNPs achieved genome-wide significance, P-value<5x10^−8^) in association with ALS. **(D)** PRS based on risk alleles of TC (independent SNPs achieved genome-wide significance, P-value<5x10^−8^) in association with ALS. LDL-C: low-density lipoprotein cholesterol; TC: total cholesterol; ALS: amyotrophic lateral sclerosis; SNPs: single nucleotide polymorphisms; log OR: log transformed odds ratio per standard deviation of PRS; SE: standard error; P_T_: P-value threshold of the association between risk alleles (SNPs) and the base phenotype; nSNP: number of independent SNPs included in the quantile. Bold black bars stand for statistically significant associations, whereas gray bars stand for associations that are not statistically significant. The gray shadows stand for the proportion of ALS variance explained by the independent SNPs in each quantile.

**Table 1.** The Best-fitted Results from the Bi-directional Polygenic Risk Score Analyses

| Base | Target | Best P<sub>T</sub> | log OR (SE) | n<sub>SNP</sub> | r<sup>2</sup> (%) | P-value |
| --- | --- | --- | --- | --- | --- | --- |
| LDL-C | ALS | 5.00x10<sup>-5</sup> | 0.15 (0.03) | 233 | 0.08 | 8.82x10<sup>-8</sup> |
| TC |  | 5.00x10<sup>-5</sup> | 0.14 (0.03) | 270 | 0.07 | 8.17x10<sup>-7</sup> |
| TG |  | 5.00x10<sup>-7</sup> | 0.07 (0.04) | 93 | 0.00 | 0.0959 |
| HDL-C |  | 0.1604 | 0.00(0.00) | 32029 | 0.00 | 0.2241 |
| LDL-C | CAD | 5.00x10<sup>-5</sup> | 0.45 (0.03) | 232 | 0.33 | 5.51x10<sup>-64</sup> |
| TC |  | 0.0125 | 0.05 (0.01) | 5094 | 0.06 | 1.36x10<sup>-12</sup> |
| TG |  | 0.0010 | 0.35 (0.02) | 832 | 0.27 | 1.36x10<sup>-53</sup> |
| HDL-C |  | 4.00x10<sup>-4</sup> | -0.29 (0.02) | 508 | 0.17 | 5.18x10<sup>-34</sup> |
| Reverse PRS analyses |  |  |  |  |  |  |
| ALS | LDL-C | 0.0009 | 0.01 (0.00) | 617 | 0.00 | 0.0859 |
|  | TC | 0.0009 | 0.01 (0.00) | 617 | 0.00 | 0.0638 |
|  | TG | 0.0001 | -0.01(0.01) | 98 | 0.00 | 0.0677 |
|  | HDL-C | 0.0001 | 0.01 (0.01) | 98 | 0.00 | 0.0772 |
| CAD | LDL-C | 0.0658 | 0.01 (0.00) | 20136 | 0.17 | 9.46x10<sup>-37</sup> |
|  | TC | 0.1478 | 0.01 (0.00) | 31666 | 0.17 | 2.14x10<sup>-39</sup> |
|  | TG | 0.0522 | 0.01 (0.00) | 17433 | 0.18 | 4.10x10<sup>-40</sup> |
|  | HDL-C | 0.0500 | -0.01 (0.00) | 17008 | 0.11 | 1.54x10<sup>-25</sup> |

In contrast, PRS based on TG and HDL-C risk alleles were not associated with ALS (Table 1). PRS based on LDL-C, TC and TG risk alleles were all statistically significantly associated with a higher risk of CAD, and PRS based on HDL-C risk alleles was associated with a lower CAD risk. In the reverse PRS analyses, PRS based on ALS risk alleles were not significantly associated with levels of any studied lipids. PRS based on CAD risk alleles were weakly but significantly associated with higher levels of LDL-C, TC, and TG whereas lower level of HDL-C.

### Causality tested by MR analyses

Based on the PRS results, we conducted MR analyses between LDL-C, TC and ALS. Because LDL-C and TC are known to be highly correlated, we chose the lead (or proxy) SNPs in all of the 27 independent loci (Supplementary Table 1) reported to be specifically associated with both LDL-C and TC, but with neither HDL-C nor TG.^13^ The combined effect of these SNPs on ALS was statistically significant (0R=1.01, 95% CI 1.00-1.02, P=0.008; Supplementary Table 2). Using these 27 SNPs as instrumental variables, the causal effects of LDL-C and TC on ALS were shown to be statistically significant (OR≥1.21, P-value≤0.001, Fig. 3A; Supplementary Fig.1A). We conducted a further analysis using 13 of the 27 SNPs, which are specific for LDL-C and TC but not associated with any other traits according to the current findings of GWAS. The causal effects of LDL-C and TC on ALS risk remained similar and statistically significant (OR≥1.26, P-value ≤0.03), and the P-value for heterogeneity became non-significant (Fig. 3B; Supplementary Fig.1B). Stronger and significant causal effects of LDL-C and TC on CAD were also identified using these sets of instrumental variables (OR⩾1.43, P-value<5x10^−5^, Supplementary Fig.2–3).

**Fig.3.**
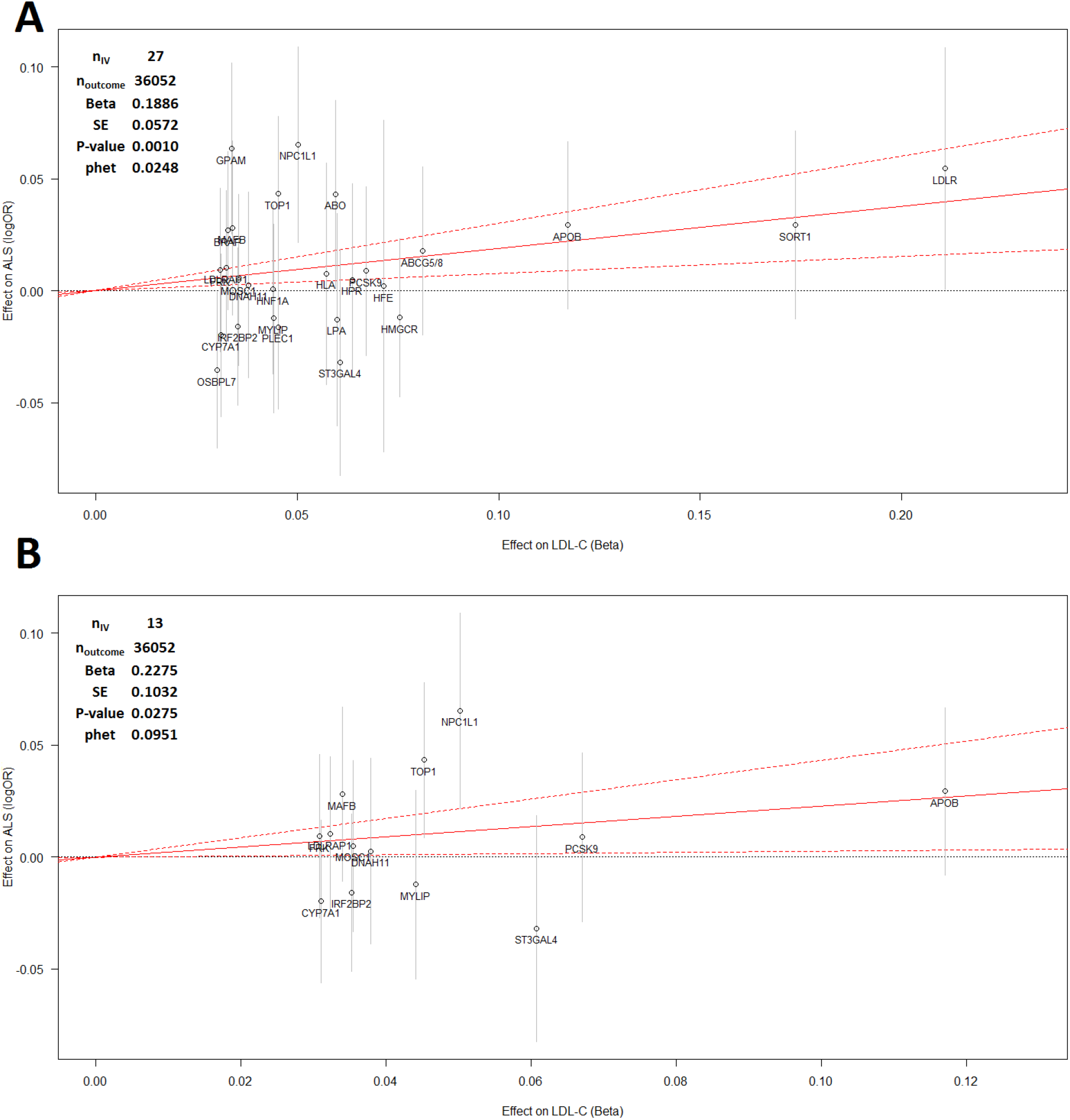
Mendelian Randomization Analysis between LDL-C and ALS in Europeans. **(A)** Instrumental variables (IVs) are lead (or proxy) SNPs in all of the 27 independent loci that are specifically associated with both LDL-C and TC (P<5xl0^−8^), but with neither HDL-C nor triglycerides (as reported by Teslovich et al., 2010). **(B)** IVs are 13 out of the 27 SNPs, which are specific for LDL-C and TC, but not associated with any other traits (screened from PhenoScanner database). The effects of IVs on ALS risk are plotted against their effects on LDL-C levels. Vertical grey lines show 95% confidence interval (95% Cl) of effect on ALS for each IV. Casual effect (log transformed odds ratio) of LDL-C on ALS risk is represented by the slope (beta) of red solid line, with red dashed lines showing the 95% Cl. n_IV_: number of IVs; n_outcome_: the sample size of genome-wide association study on ALS. phet: the P-value of heterogeneity test. SNPs: single nucleotide polymorphisms; LDL-C and HDL-C: low- and high- density lipoprotein cholesterol; TC: total cholesterol; ALS: amyotrophic lateral sclerosis.

## Discussion

This is the first study using polygenic methods on summary GWAS results to test the association and causality between blood lipids and ALS. When using unrestricted but independent SNPs across the genome, we find PRS based on LDL-C and TC risk alleles to be significantly associated with ALS risk. When using restricted and independent SNPs that are specifically associated with LDL-C and TC but not with other lipids or risk factors in the MR analysis setting, we further identified evidence for a causal relationship between LDL-C, TC and ALS.

In the bi-directional PRS analyses, we found that PRS based on LDL and TC risk alleles were significantly associated with ALS risk, whereas PRS based on ALS risk alleles were not associated with any of the studied lipids. One major reason might be the smaller sample size of ALS GWAS compared to GWAS of blood lipids and CAD, which would influence the precision of the estimated effect sizes and thereby their predictive ability. Moreover, SNPs in the best-fitted quantiles explain extremely low variance (close to zero) of the target, whereas in the analysis of lipids and CAD, SNPs in the best-fitted quantiles explain 0.1∼0.3% of the target in both direction (Table 1). A first explanation is similarly the inadequate sample size of ALS GWAS, with less than eight genome-wide significant loci have been identified to be associated with ALS in the latest study.^14^ Another explanation might be the special polygenic rare variant architecture of ALS. Unlike most common complex diseases, low-frequency or rare alleles may explain proportionally more heritability of ALS.^14^ Because GWAS only capture contributions from common SNPs with minor allele frequency over 1% in the population, it is insensitive to capture effects from rare variants. This explain partly the reason why SNP-based heritability of ALS estimated from GWAS was less than 9%, whereas previous family and twin studies reported a heritability of ALS as high as 60%.^19^ Such a possibility is further supported by the fact that the estimates of OR and explained variance of ALS stay at a stable level from P_T_=5x10^−19^ to 5x10^−5^, but become smaller and not statistically significant as additional and less informative or less specific SNPs (P_T_>5x10^−5^) were included (Fig. 2).

In the MR analyses, we identified statistically significant causal effects of LDL-C and TC on ALS, when using specific genetic variants for LDL-C and TC as the instrumental variables. These genetic variants are in the loci close to the genes involved in the cholesterol transportation or lipid modification processes, which may point to potential polygenic pathways between dyslipidemia and ALS (Supplementary Table 2).

Limitations of the presented polygenic methods should be acknowledged. First, in the PRS analyses, we grouped independent SNPs (by LD clumping) into 5000 quantiles with gradually increasing P-value thresholds (Pt from 0 to 0.5) across the whole genome. The threshold that explains the largest variance was defined as the best-fitted Pt. Such optimization over a particular parameter may introduce some degree of over-fitting (winners curse). However, even if we did apply overly conservative Bonferroni correction with a new significance level at 1x10^−5^, our findings would still remain statistically significant, arguing against the possibility that our conclusion is sensitive to such potential over-fitting. Second, we should bear in mind that the causality tested by MR analysis is based on certain assumptions. The most important one is that instrumental variables should be specific for exposure but not associated with other confounding factors.^12^ To test the potential influence of this assumption, we used PhenoScanner to further select 13 SNPs that are specific for LDL-C and TC and not associated with any other traits according to current GWAS findings. As the results are largely similar (Fig.3; Supplementary Fig.1–3), the concern about MR assumption was relieved.

Although our study found clear polygenic evidence that LDL-C and TC are causally associated with ALS, larger GWAS and deep sequencing for rare variants might be essential to further understand the genetic mechanisms of dyslipidemia in ALS. In parallel, functional molecular studies are needed to shed light on potential pathogenic mechanisms of the suggested polygenic pathways relating lipid processing to ALS risk.

## Disclosure statement

The authors have no conflicts of interest to disclose.

## Acknowledgements

We would like to thank the following consortia for sharing their summary GWAS results: the Global Lipids Genetics Consortium, CARDIoGRAMplusC4D Consortium and Project MinE GWAS Consortium. This study was funded by the Swedish Research Council (grant no. 2015-03170), the Karolinska Institutet (Senior Researcher Award and Strategic Research Program in Epidemiology), the Ulla-Carin Lindquist Foundation, and the China Scholarship Council (grant no. 201306210065).

## Web Resources

Summary GWAS results of blood lipids, http://csg.sph.umich.edu/abecasis/public/

Summary GWAS result of ALS, http://databrowser.projectmine.com/

Summary GWAS results of CAD, http://www.cardiogramplusc4d.org/data-downloads/

The HapMap reference genotype, http://zzz.bwh.harvard.edu/plink/res.shtml#hapmap

PRSice: Polygenic Risk Score software, http://prsice.info/

PhenoScanner, http://www.phenoscanner.medschl.cam.ac.uk

GTX: Genetics ToolboX, https://cran.r-project.org/web/packages/gtx/

